# Subcellular Localization Constrains Protein Detectability and Reveals Systematic RNA-Protein Discordance Across Cancers

**DOI:** 10.64898/2026.03.30.713919

**Authors:** Kedar Joshi, Saniya Kate

**Affiliations:** Omnistrata AI; Metaphore

## Abstract

Transcript abundance is widely used as a proxy for protein expression in cancer studies; however, mRNA levels often fail to predict protein detectability due to post-transcriptional and compartment-specific regulatory processes. Here, we present a machine learning framework that integrates RNA expression, gene-level attributes, and subcellular localization to model protein detectability across human cancers.

Leveraging transcriptomic data from TCGA, TARGET, and GTEx, and protein annotations from the Human Protein Atlas, we constructed a dataset comprising over 100,000 gene–cancer pairs across seven tumor types. Models based on RNA features alone achieved moderate predictive performance (ROC-AUC ~0.71), whereas incorporating subcellular localization significantly improved accuracy (ROC-AUC ~0.82). Paired bootstrap analysis confirmed that these gains were statistically robust.

We further identify a substantial set of genes with high transcript abundance yet absent protein detection, revealing widespread RNA-protein decoupling. These discordant genes are enriched in mitochondrial, metabolic, and translational regulatory pathways, suggesting that discordance reflects structured biological processes rather than stochastic variation. Together, our results demonstrate that cellular context, particularly subcellular localization, is a key determinant of protein detectability and underscore the limitations of transcript-centric interpretations in cancer genomics.

## Introduction

High-throughput transcriptomic profiling has become central to cancer research, with RNA abundance frequently used as a surrogate for protein expression. However, numerous studies have demonstrated that mRNA levels explain only a fraction of the variability observed at the protein level (Liu et al., 2016; Vogel & Marcotte, 2012). This discrepancy arises from multiple layers of regulation, including translation efficiency, protein degradation, and spatial compartmentalization within the cell.

Despite these limitations, RNA-seq remains the dominant modality in large-scale cancer studies, and systematic approaches to model RNA-protein relationships across tumor types remain underdeveloped. In particular, the extent to which biological context such as subcellular localization, constrains protein detectability has not been quantitatively assessed at scale.

Here, we develop a predictive framework that integrates transcriptomic features with gene-level attributes and subcellular localization to model protein detectability across cancers. By combining data from large consortia with curated protein annotations, we aim to (i) quantify the predictive limits of RNA expression, (ii) evaluate the contribution of biological context, and (iii) systematically characterize genes exhibiting RNA–protein discordance.

To our knowledge, this study provides one of the first large-scale, cross-cancer analyses linking subcellular localization to protein detectability and formally quantifying RNA–protein decoupling using a predictive modeling approach.

## Methods

### Data sources and preprocessing

RNA expression data were obtained from the UCSC Xena recompute compendium (TCGA, TARGET, and GTEx). Protein detectability and subcellular localization annotations were derived from the Human Protein Atlas (HPA) (Uhlén et al., 2015; Thul et al., 2017).

Gene expression was summarized at the gene–cancer level using log2-transformed TPM values and tumor–normal fold change. Gene identifiers were harmonized by converting to uppercase gene symbols prior to merging across datasets. Subcellular localization annotations were encoded as binary indicators, with missing entries set to zero following integration.

The final dataset comprised over 100,000 gene–cancer pairs across seven cancer types (BRCA, COAD, GBM, LIHC, PAAD, PRAD, THCA).

Missing feature values were handled using median imputation within model pipelines.

### Feature construction

Predictive features included tumor RNA expression (log2 TPM mean), tumor–normal RNA fold change, gene length, protein-coding status, and subcellular localization indicators derived from HPA annotations.

Model inputs were defined as follows:

- RNA-only model: RNA expression and fold change
- RNA + gene-level model: RNA features plus gene length and coding status
- Localization-aware model: RNA features, gene length, and subcellular localization

### Protein detectability definition

Protein detectability was defined as a binary outcome based on aggregated HPA immunohistochemistry data. A gene was considered protein-detectable if the number of samples with reported expression (high, medium, or low) exceeded those with no detection, providing a conservative estimate of protein presence.

### Modeling framework

Logistic regression models were implemented using scikit-learn pipelines consisting of median imputation, feature standardization, and class-balanced logistic regression (max_iter = 1000).

Random forest models were implemented using 400 trees with class-balanced weighting, random_state = 7, and parallel computation enabled. No hyperparameter tuning was performed, and all models were evaluated using fixed configurations.

Feature importance was derived from the fitted random forest model

### Cross-cancer evaluation

Model performance was assessed using a leave one cancer out (LOCO) framework, in which each cancer type was held out as an independent test set. This design evaluates generalization across biological contexts rather than within-cohort performance.

Predictive performance was quantified using ROC-AUC and PR-AUC. Confidence intervals were estimated using 1,000 bootstrap resamples of test predictions.

### Statistical comparison of models

To compare models on identical samples, paired bootstrap analysis was performed using 5,000 resamples. Differences in ROC-AUC and PR-AUC were summarized using empirical confidence intervals and two-sided p-values.

### Identification of RNA–protein discordance

RNA–protein discordance was defined as genes with high RNA expression but no detectable protein. High expression was defined using quantile-based thresholds within each cancer type. Using out-of-fold predictions from the localization-aware model, discordant candidate genes were identified as those with:

- high RNA expression
- high predicted probability of protein detectability
- observed absence of protein detection

Sensitivity analyses were conducted across multiple RNA quantile thresholds (0.75, 0.80, 0.90) and prediction probability thresholds (0.70, 0.80, 0.90).

## Results

Model performance and downstream analyses are summarized in Figures 1–5. Subcellular

**Figure 1.**
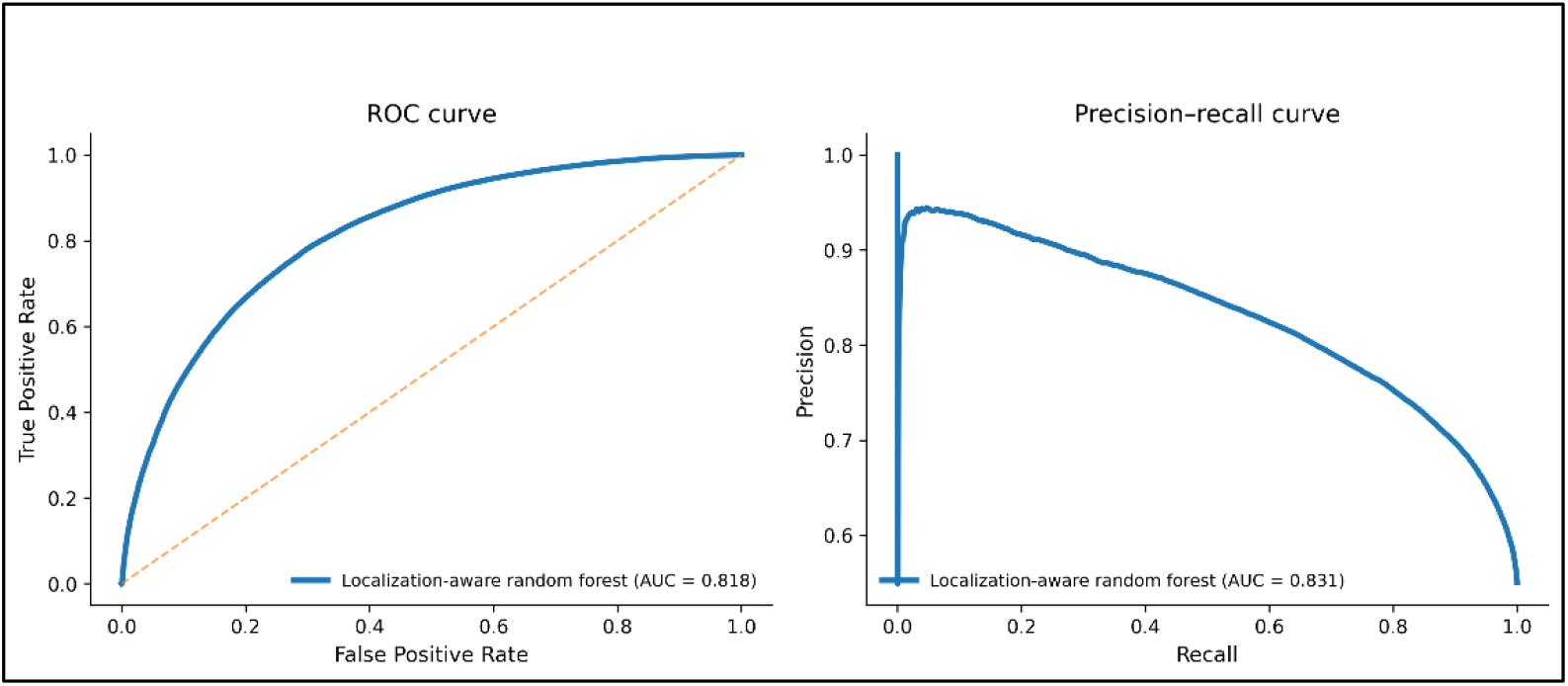
Subcellular localization substantially improves prediction of protein detectability.

**Figure 2.**
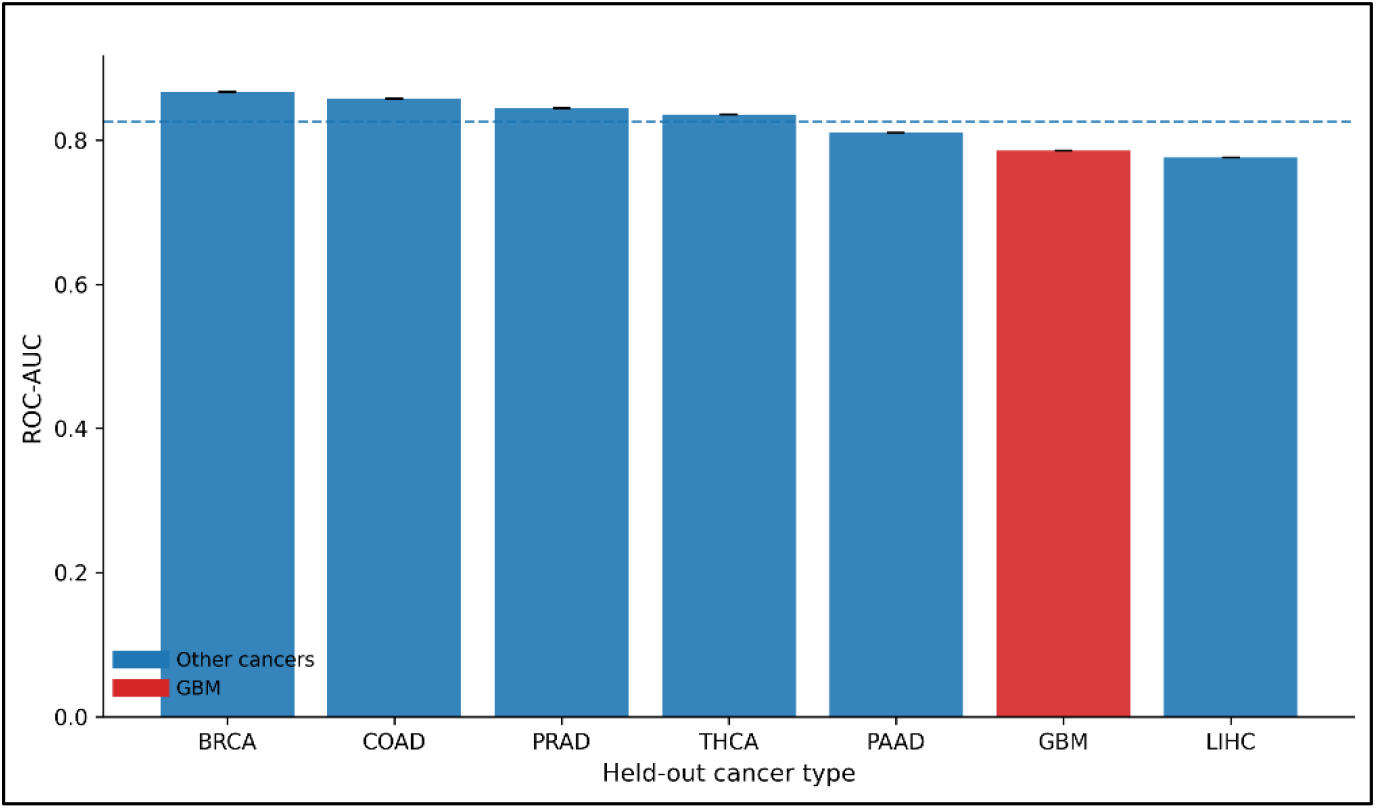
Generalization of protein detectability prediction across cancer types.

**Figure 3.**
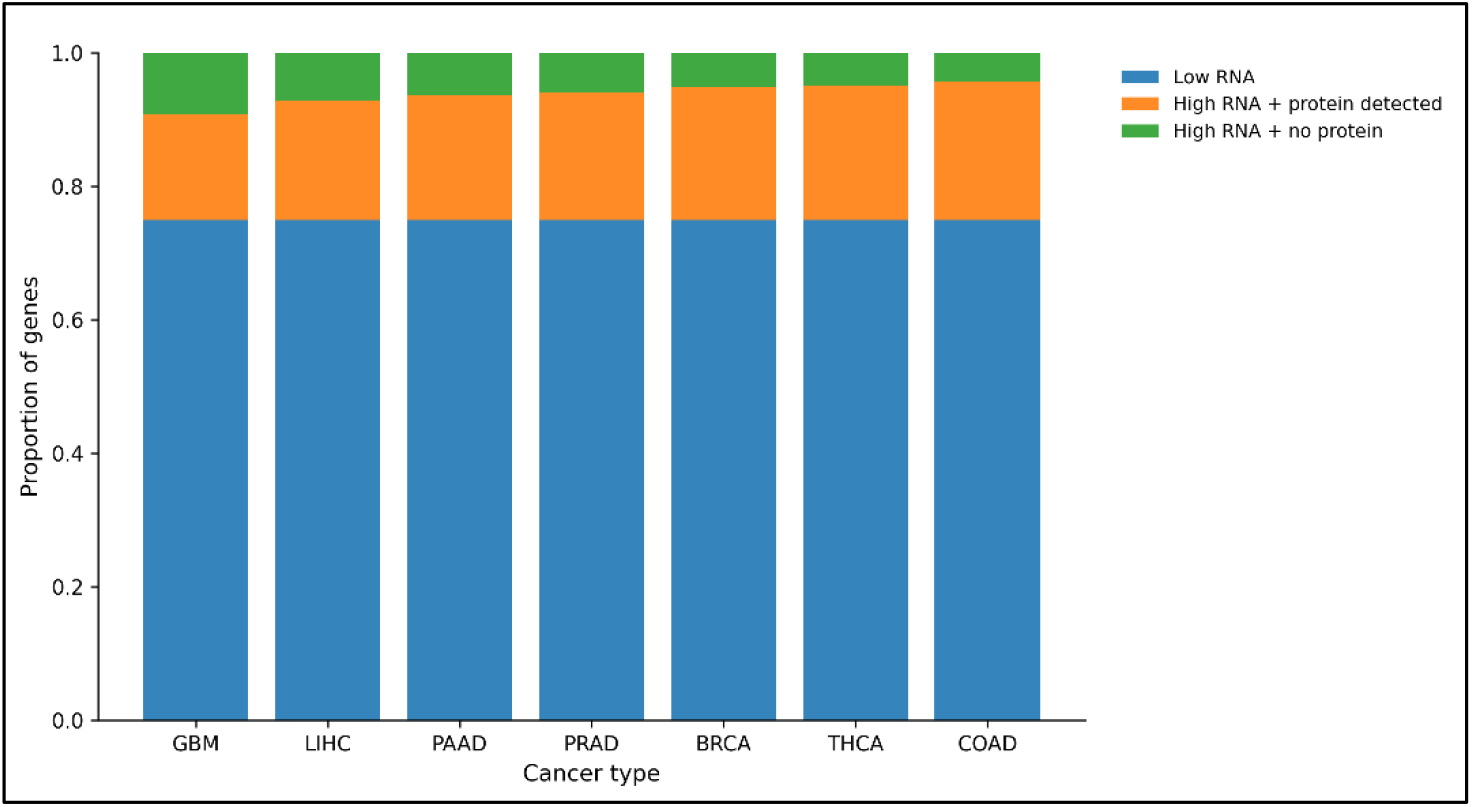
Widespread RNA–protein discordance across cancer types.

**Figure 4.**
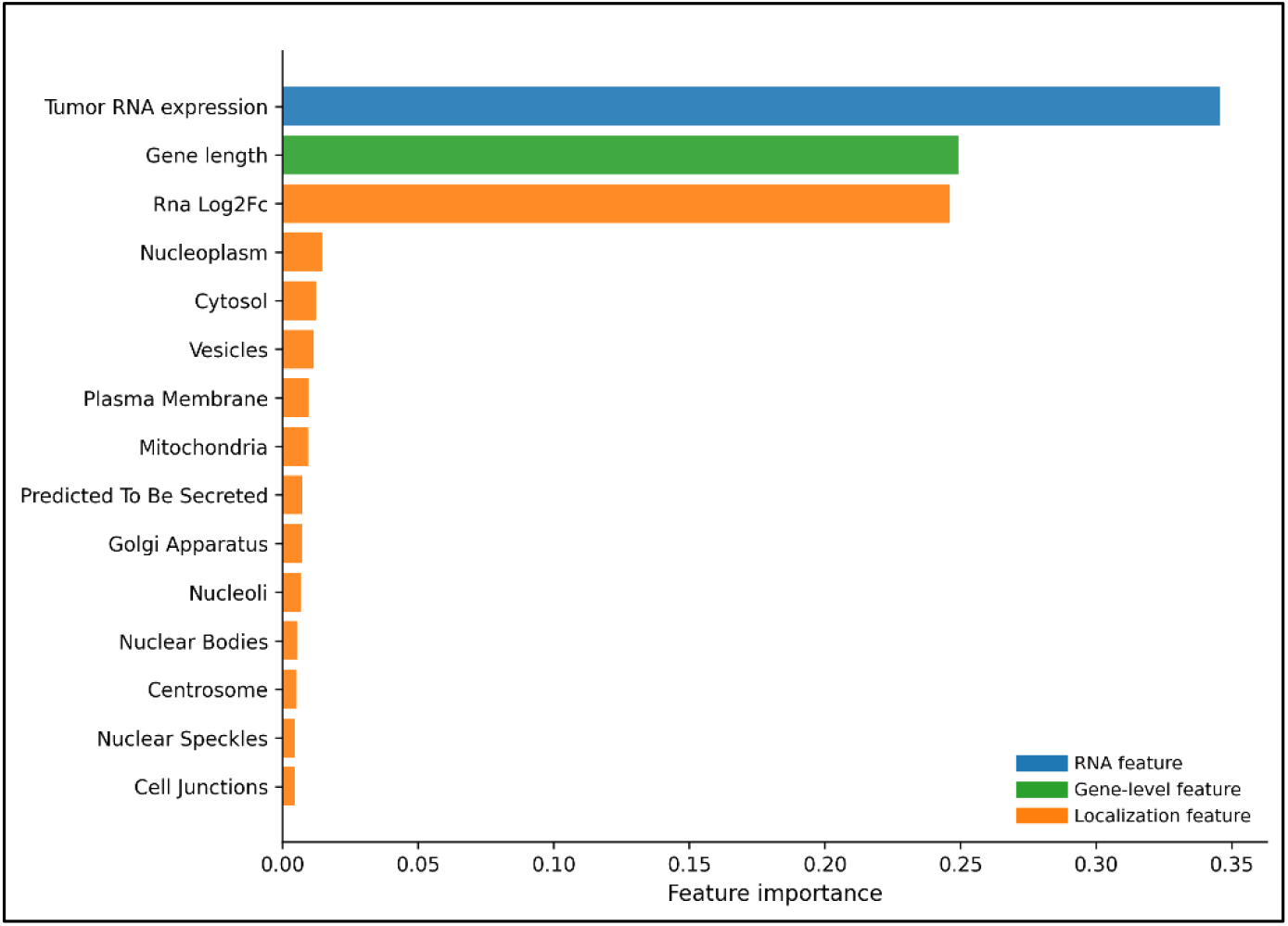
Feature importance highlights the role of subcellular localization.

**Figure 5.**
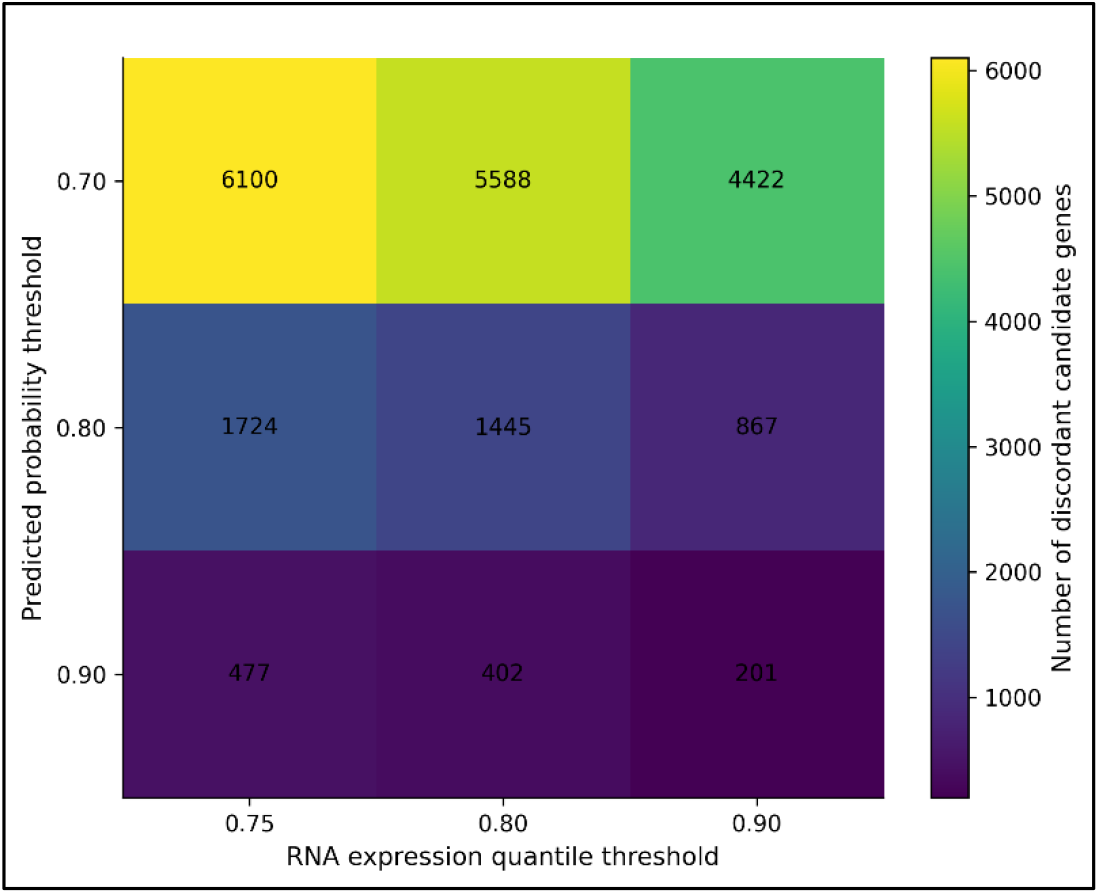
Robustness of RNA–protein discordance across threshold definitions.

### localization improves prediction of protein detectability

Models based on RNA expression alone achieved moderate performance (ROC-AUC ~0.71). Incorporation of gene-level features provided limited additional gain. In contrast, the localization-aware model substantially improved predictive performance (ROC-AUC ~0.82), with paired bootstrap analysis confirming statistically significant improvements over RNA-only models. These results indicate that subcellular localization encodes critical biological constraints on protein detectability that are not captured by transcript abundance.

### Generalization across cancer types

Performance remained consistent across cancer types under leave-one-cancer-out evaluation. However, glioblastoma exhibited reduced predictive accuracy relative to other cancers, suggesting increased regulatory complexity and a greater degree of RNA–protein decoupling in this tumor type.

### Widespread RNA–protein discordance across cancers

Across all cancer types, a substantial fraction of genes exhibited discordance between RNA expression and protein detectability. This discordance varied across cancers, with glioblastoma showing the highest rates, consistent with enhanced post-transcriptional regulation. These findings indicate that the relationship between transcript abundance and protein detectability is context-dependent and influenced by tumor-specific regulatory mechanisms.

### Biological characterization of discordant candidate genes

Discordant candidate genes were enriched in specific biological categories, including mitochondrial proteins (MRPL15, COX6A1, COX6C), metabolic enzymes (ALDOA), and RNA-binding proteins (CSDE1). These patterns suggest that RNA–protein discordance reflects structured regulatory processes, particularly those involving mitochondrial function, metabolic adaptation, and translational control.

Receiver operating characteristic (ROC) curves (left) and precision–recall (PR) curves (right) comparing models with increasing biological context: RNA-only logistic regression, RNA plus gene-level features, and a localization-aware random forest model. Performance is evaluated using leave-one-cancer-out cross-validation. The localization-aware model consistently outperforms RNA-based models, demonstrating that subcellular localization provides critical information beyond transcript abundance for predicting protein detectability. Reported curves represent aggregated out-of-fold predictions across all cancer types.

Model performance (ROC-AUC) for the localization-aware model across individual cancer types under leave-one-cancer-out evaluation. Each bar represents performance on a held-out cancer type. Performance remains consistently high across cancers, indicating robust generalization. Reduced accuracy in glioblastoma (GBM) suggests increased regulatory complexity and a greater degree of RNA–protein decoupling in this tumor type.

Proportion of genes exhibiting RNA–protein discordance across cancer types, defined as genes with high RNA expression (top quartile) but no detectable protein. Discordance rates vary across cancers, with glioblastoma showing the highest levels. These findings indicate that the relationship between transcript abundance and protein detectability is context-dependent and influenced by tumor-specific regulatory mechanisms.

Feature importance derived from the localization-aware random forest model. Both RNA expression features and subcellular localization variables contribute substantially to model performance. Localization features rank among the most informative predictors, supporting the role of cellular compartmentalization in constraining protein detectability.

Sensitivity analysis shows the number of discordant candidate genes identified across combinations of RNA expression thresholds and predicted probability thresholds. While absolute counts vary with threshold stringency, the presence of a substantial set of discordant genes is consistent across parameter choices, indicating that the observed RNA–protein decoupling is robust.

## Discussion

In this study, we systematically evaluated the extent to which RNA expression predicts protein detectability across cancer types and shows that transcript abundance alone provides limited predictive power. Incorporating subcellular localization substantially improves model performance, demonstrating that cellular context plays a critical role in constraining protein detectability.

These findings have important implications for the interpretation of transcriptomic data in cancer research. RNA expression is frequently used as a surrogate for protein abundance in biomarker discovery and pathway analysis; however, our results indicate that this assumption can be unreliable for a substantial subset of genes. Predictive frameworks that incorporate biological context provide a more accurate representation of functional protein output.

The strong contribution of subcellular localization suggests that compartment-specific regulatory processes influence whether proteins are detectable. Proteins localized to different cellular regions are subject to distinct mechanisms, including compartment-specific translation, protein trafficking, and degradation. These processes are not captured by transcript abundance alone but are partially reflected through localization features, explaining their impact on predictive performance.

We also identify widespread RNA–protein discordance across cancers, with many genes exhibiting high RNA expression but no detectable protein. Importantly, these genes are enriched in specific functional categories, indicating that discordance reflects structured biological regulation rather than random variation. Mitochondrial proteins, metabolic enzymes, and RNA-binding proteins are particularly prominent among discordant candidates.

These functional patterns point to multiple mechanisms underlying RNA–protein decoupling. Mitochondrial proteins are governed by distinct translational systems and import processes, which may uncouple transcript abundance from detectable protein levels. Metabolic enzymes operate within dynamically regulated pathways in cancer, where transcriptional changes do not necessarily translate to protein-level changes. The presence of RNA-binding proteins further implicates post-transcriptional control, as these factors regulate mRNA stability and translation efficiency independently of transcript abundance.

Variation in predictive performance across cancer types further supports a context-dependent relationship between RNA and protein expression. Glioblastoma shows reduced model accuracy and elevated discordance, consistent with its known alterations in translational regulation and protein turnover. This highlights the influence of tumor-specific regulatory programs on RNA–protein relationships.

Technical factors may also contribute to the observed discordance. Protein detectability in the Human Protein Atlas is based on immunohistochemistry, which depends on antibody specificity, epitope accessibility, and tissue context. These constraints may lead to under-detection of certain proteins, particularly those associated with membranes or structurally complex compartments.

Several limitations should be considered. RNA and protein data were derived from independent datasets and are not matched at the sample level, which may introduce variability. Protein detectability was defined as a binary outcome, simplifying a continuous biological process.

Additionally, the identification of discordant genes relies on threshold-based definitions, although sensitivity analyses indicate that the overall patterns are robust. Differences in data generation protocols may also introduce batch effects that are not fully accounted for.

Despite these limitations, this study provides a scalable framework for modeling RNA–protein relationships across cancers. By integrating transcriptomic features with biological context, we show that protein detectability is shaped by both expression and cellular localization, and that RNA–protein discordance reflects structured regulatory processes.

## Conclusion

In summary, RNA expression alone provides limited predictive power for protein detectability across cancers, while incorporation of subcellular localization substantially improves model performance. We identify widespread and biologically structured RNA–protein discordance, particularly among genes involved in mitochondrial function, metabolism, and translational regulation. These findings highlight the importance of incorporating biological context into multi-omics analyses and caution against transcript-centric interpretations in cancer research.

## Data Availability

All datasets used in this study are publicly available from the UCSC Xena platform (TCGA, TARGET, GTEx) and the Human Protein Atlas.

## Code Availability

Analysis code was implemented in Python using pandas, NumPy, scikit-learn, and joblib. Scripts for model training, cross-cancer evaluation, statistical comparison, and discordance analysis will be made publicly available on GitHub upon publication.

## References

1. Liu, Y., Beyer, A., & Aebersold, R. (2016). On the dependency of cellular protein levels on mRNA abundance. Cell, 165(3), 535–550. 10.1016/j.cell.2016.03.014

2. Vogel, C., & Marcotte, E. M. (2012). Insights into the regulation of protein abundance from proteomic and transcriptomic analyses. Nature Reviews Genetics, 13(4), 227–232. 10.1038/nrg3185

3. Thul, P. J., Åkesson, L., Wiking, M., Mahdessian, D., Geladaki, A., Ait Blal, H., Alm, T., Asplund, A., Björk, L., Breckels, L. M., Bäckström, A., Danielsson, F., Fagerberg, L., Fall, J., Gatto, L., Gnann, C., Hober, S., Hjelmare, M., Johansson, F., … Uhlén, M. (2017). A subcellular map of the human proteome. Science, 356(6340), eaal3321. 10.1126/science.aal3321

4. Sonenberg, N., & Hinnebusch, A. G. (2009). Regulation of translation initiation in eukaryotes: Mechanisms and biological targets. Cell, 136(4), 731–745. 10.1016/j.cell.2009.01.042

5. Liberti, M. V., & Locasale, J. W. (2016). The Warburg effect: How does it benefit cancer cells? Trends in Biochemical Sciences, 41(3), 211–218. 10.1016/j.tibs.2015.12.001

6. Uhlén, M., Fagerberg, L., Hallström, B. M., Lindskog, C., Oksvold, P., Mardinoglu, A., Sivertsson, Å., Kampf, C., Sjöstedt, E., Asplund, A., Olsson, I., Edlund, K., Lundberg, E., Navani, S., Szigyarto, C. A. K., Odeberg, J., Djureinovic, D., Takanen, J. O., Hober, S.,… Pontén, F. (2015). Tissue-based map of the human proteome. Science, 347(6220), 1260419. 10.1126/science.1260419

